# A Novel Algorithm to Calculate Elementary Modes: Analysis of *Campylobacter jejuni* Metabolism

**DOI:** 10.1101/2023.01.11.521685

**Authors:** Yanica Said, Dipali Singh, Cristiana Sebu, Mark Poolman

## Abstract

We describe a novel algorithm, ‘LPEM’, that given a steady-state flux vector from a (possibly genome-scale) metabolic model, decomposes that vector into a set of weighted elementary modes such that the sum of these elementary modes is equal to the original flux vector.

We apply the algorithm to a genome scale metabolic model of the human pathogen *Campylobacter jejuni*. This organism is unusual in that it has an absolute growth requirement for oxygen, despite being able to operate the electron transport chain anaerobically.

We conclude that 1) Microaerophilly in *C. jejuni* can be explained by the dependence of pyridoxine 5′ -phosphate oxidase for the synthesis of pyridoxal 5′ - phosphate (the biologically active form of vitamin B6), 2) The LPEM algorithm is capable of determining the elementary modes of a linear programming solution describing the simultaneous production of 51 biomass precursors, 3) Elementary modes for the production of individual biomass precursors are significantly more complex when all others are produced simultaneously than those for the same product in isolation and 4) The sum of elementary modes for the production of all precursors in isolation requires a greater number of reactions and overall total flux than the simultaneous production of all precursors.

## 1 Introduction

### 1.1 Camplylobacter

Campylobacter, a genus of Gram-negative, curved or spiral, highly motile bacilli, are foodborne pathogens and the leading cause of acute bacterial gastroenteritis worldwide. The most common routes of human infection is fecal-oral or via contaminated meat, with poultry, in which it is a common gut commensal, being of particular concern. However, it is also found in the gut of wild birds, other farmed animals such as pigs and cattle, pets, as well as soil and contaminated water sources. The ability of this pathogen to persist in the food chain and to contaminate food products poses a serious challenge for food safety and global health.

Campylobacteriosis presents with typical symptoms of gastrointestinal infection, including diarrhoea (often bloody), fever, nausea, and abdominal pain. It is a notifiable disease in the UK with approximately 52,000 cases reported in 2016^1^, with an annual cost estimated as _£_700 million Daniel et al. [2020]. Although the disease is commonly self-limiting, it can be associated with a number of serious or life-threatening sequelae including colitis, reactive arthritis, and Guillain-Barré syndrome (GBS), a neurological autoimmune disorder causing muscle weakness and even paralysis requiring ventilation in severe cases. The association of *Campylobacter spp*. with GBS is assumed to be due to the similarity of a lipopolysaccharide cell wall component with a human ganglioside Hadden et al. [2002], Poropatich et al. [2010].

#### 1.1.1 Microbiology and biochemistry

*Campylobacter jejuni* is well known for being microaerophilic, having an absolute requirement for oxygen and growing optimally at a pO_2_ of *≈* 50 mbar but is non-viable (or at least unculturable) at atmospheric pO_2_. The reason for the requirement for O_2_ is not well understood, although as *C. jejuni* is able to respire anaerobically, it has been proposed that this is a biosynthetic requirement. It is of note that although the intestinal lumen is generally thought of as an anaerobic environment, in regions close to the mucosa, pO_2_ has been reported as being as high as 60 mbar Albenberg et al. [2014], Zheng et al. [2015].

Another unusual feature of *C. jejuni* metabolism is that it lacks the enzymes forming the initial steps of glycolysis (glucokinase and phosphofructokinase) as well as those of oxidative limb of the oxidative pentose phosphate pathway (glucose-6-phosphate dehydrogenase, phosphoglu-conolactonase and phosphogluconate dehydrogenase) Parkhill et al. [2000], Stahl et al. [2012]. It also lacks the monosaccharide transporters found in most other bacteria, although it has been suggested that some *Campylobacter spp*. strains are capable of catabolising fucose (a common saccharide residue in intestinal mucins), otherwise the preferred carbon sources are pyruvate, TCA cycle intermediates, and amino acids Hofreuter [2014], Tejera et al. [2020], Wagley et al. [2014].

*C. jejuni* possess a branched electron transport chain (ETC) with the flexibility to utilise a number of substrates as electron donors including H_2_ and formate (both present in the intestinal lumen as metabolic by-products of other gut microbes) Bernalier-Donadille [2010], Kelly [2008], Weerakoon et al. [2009]. O_2_ can act as the terminal electron acceptor, although NO_3_, NO_2_, SO_4_ and fumarate can also fulfil this function Stoakes et al. [2022], van der Stel and Wösten [2019], van der Stel et al. [2017], allowing for the anaerobic operation of the ETC. However, despite this, *C. jejuni* is unable to grow anaerobically Kaakoush et al. [2007], Sellars et al. [2002] and this has been proposed to be due the existence of oxygen dependent reactions involved in the synthesis of one or more biomass components, specifically: 1) coproporphyrinogen oxidase (EC 1.3.3.3) - a component of heme biosynthesis, and ii) that O_2_ is required for the post-translational modification of the enzyme ribonucleotide reductase (RNR) Sellars et al. [2002]. However, as it has also been reported that the *C. jejuni* genome encodes for the O_2_ independent coproporphyrinogen dehydrogenase (EC 1.3.98.3) capable of fulfilling the same physiological role as its O_2_ dependent counterpart de Vries et al. [2015], Parkhill et al. [2000], in which case the latter proposal would appear unlikely to be correct.

In order to further investigate the O_2_ dependence of *Campylobacter spp*. we here describe the analysis of a genome-scale metabolic model (GSM) of *C. jejuni* in order to investigate oxygen utilisation for the production of individual biomass precursors whilst growing in a minimal medium. We also present a novel algorithm to identify the pathways (Elementary Modes - see below) utilised when all biomass precursors are synthesised simultaneously.

### 1.2 Modelling Background

GSMs are computational representations of all reactions (assumed to be) present in an organism’s repertoire, typically derived in the first instance from an annotated genome, with reactions defined in terms of stoichiometry, directionality and reversibility, but without any information about reaction kinetics. Such models can be referred to as stoichiometric, or *structural* models.

The theoretical basis for the analysis of structural models can be divided into two broad categories, those derived from *null-space* analysis, and in particular Elementary Modes Analysis (EMA) Schuster et al. [1999, 2000], and those derived from *linear programming* (LP) Fell and Small [1986], Orth et al. [2010]; both assume steady-state conditions and both can be regarded as pathway identification tools. The essential difference between the two is that whilst EMA identifies all possible independent routes through the network, LP results typically identify a single (possibly non-unique) pathway through the network that satisfies given optimisation criteria, and the two can be regarded as complementary approaches.

An Elementary Mode (EM) is a minimal metabolic sub-network capable of sustaining a steady-state flux whilst respecting reversibility criteria, associated with a net conversion of substrates into products, and any steady-state flux distribution of the network can be obtained from a non-negative summation of EMs Schuster and Hilgetag [1994], Schuster et al. [2000]. Given the complete set of EMs for a network and an associated flux vector, it is possible to then calculate the flux carried by each EM Poolman et al. [2004], Schwartz and Kanehisa [2005].

A well-known draw-back of EMA is the fact that it suffers from combinatorial explosion (is NP-hard - Acuña et al. [2009], Klamt and Gilles [2004]), and therefore obtaining the complete set of EMs of a GSM is impractical for most purposes, not only in terms of the computational resources required but also the unwieldy size of any resultant data-set should such a calculation be possible.

In contrast, flux vectors (modes which may or may not be elementary) with defined properties can be obtained very rapidly using LP; methods based on this (such those related to flux balance analysis (FBA) Orth et al. [2010], Schilling et al. [2000]) are currently the predominant means of analysing GSMs. One draw-back of using LP in the context of a GSM, and in particular where the aim of the study is to account for growth (and therefore the production of all biomass precursors), is that the resulting flux vector still defines a large network that requires further analysis to be easily understood.

However, the solution to a given linear program applied to a GSM will generally contain a much smaller number of reactions than the original, and under some circumstances may be small enough to be treated as a sub-network amenable for EMA [Hartman et al., 2014, Mesfin, 2020].

A number of methods for decomposing flux-vectors into EMs without having previously determined the complete set of EMs have also been described. These utilise LP and/or Mixed Integer Linear Programming (MILP) as strategies for finding individual EMs that fulfil a set of desired attributes. Oddsdóttir et al. [2015], obtained EMs that account for a set of observed transporter fluxes. Their algorithm starts from a matrix, **E**, whose columns, **E**_*i*_, comprise of a small sub-set of EMs, and a weighting vector, **w**. Two optimization problems are simultaneously solved; a least-squares data fitting master problem that iteratively improves upon the weighting vector **w**, aiming to obtain a product, **Ew**, that is consistent with the external flux measurements, and an additional sub-problem that after each iteration of the master, obtains a new EM to be added to the matrix **E** such that the least-square fit is improved. The algorithm concludes when the fit can no longer be improved by the addition of more EMs. This strategy requires solving a quadratic programme at each iteration, which due to being inherently more computationally costly than LP limits this algorithm’s suitability for large models.

Hung et al. [2011], proposed an algorithm that decomposes a steady-state flux vector, **v**, in series of stages. At each stage, a succession of MILPs reduce the number of non-zero fluxes in **v** (whilst considering the values of **v** as an upper-bound) until a solution which cannot be reduced further (and therefore is an EM) is obtained. The EM is subtracted from **v** and the process is repeated until all constituent EMs have been discovered. A drawback of this algorithm is the heavy use of MILPs, which are more computationally costly then LP.

Jungers et al. [2011], used LP to decompose flux vectors into a minimal number of EMs. This algorithm extracts EMs from the solution-space generated by combining the network’s stoichiometry with the steady-state flux-vector, **v**. The first EM is chosen at random from this space, whilst subsequent EMs are chosen such that the difference from them and the previous modes is maximized. This technique reduces the dimension of the solution-space one step at a time, and, as a consequence has the advantage that the maximum number of EMs obtained must be equivalent to the dimension of the null-space. However, it is likely to produce different results each time that it is used, making the replication of findings difficult.

In this work we describe an algorithm (LPEM) which uses LP to extract a set of EMs that account for a steady-state flux vector (which itself may be an LP solution) from a (possibly genome-scale) metabolic model and demonstrate its application and utility by investigating the phenomenon of micro-aerophilly in *C. jejuni*. The approach is simialr to that described by Hung et al. [2011] but has the advantage that it requires only LP, and not MILP. This algorithm is implemented as part of the open-source metabolic modelling software package, ScrumPy Poolman [2006, 2020].

## 2 Methods

### 2.1 The Model

The model used here is that described by Stoakes et al. [2022] (a revision of that previously described by Tejera et al. [2020]). As the genome lacks annotation for enzymes involved in glyoxylate shunt (malate synthase and isocitrate lyase) there is no sink for glycolaldehyde, (a by-product of folate synthesis) and therefore a corresponding transporter added. The model consists of 1029 reactions, 92 transporters and 988 metabolites. It is capable of producing biomass components in a defined media as described in Tejera et al. [2020].

### 2.2 Model analysis

Initial model analysis was performed using the linear program defined by Eq. (1).

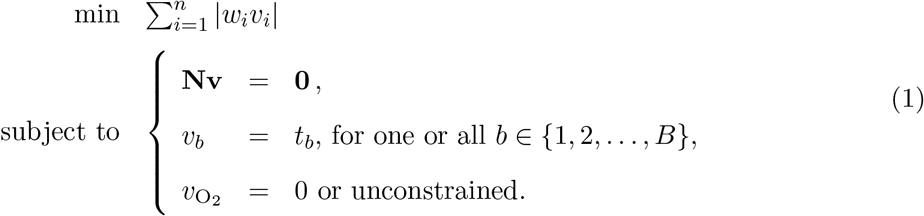

The objective is to determine a flux vector **v** whose weighted sum of fluxes is minimised. Each component, *v*_*i*_, *i* = (1…*n*), where *n* is the number of reactions, *v*_*i*_ corresponds to the flux being carried by the *i*^th^ reaction and *w*_*i*_ is the associated weighting coefficient. Except where specified, weighting coefficients, *w*_*i*_, were set to 1 giving equal weight to the minimisation of flux carried by each reaction.

This is subject to the steady-state condition **Nv** = **0** and the production of one or more biomass components may be taken into account by setting a fixed constraint, *v*_*b*_ = *t*_*b*_, where the suffix *b* denotes the *b*^*th*^ out of *B* exported biomass components whose transport flux, *v*_*b*_, is defined by its relative abundance, *t*_*b*_ (m.gdw^*−*1^) multiplied by the growth-rate which, for the purposes of this study, was arbitrarily set to unity. In order to investigate the O_2_ requirement the associated transport flux, 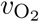, was either set to zero or unconstrained and the corresponding penalty weighting, 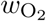, was set either to 1 or 10^6^ as described below.

With 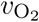 set to zero, in combination with a demand for all biomass components, Eq. (1) has no feasible solution, demonstrating that at least one biomass component has an absolute requirement for O_2_. In order to identify the O_2_ dependent biomass component(s), Eq. (1) was solved repeatedly for each biomass component at a time, with the oxygen transport flux constrained to zero. Obtaining a feasible solution with such a constraint demonstrates that this component can be produced without oxygen, conversely, failure to obtain a solution demonstrates that the synthesis of that component has an absolute dependence on oxygen.

**Panel 1.**
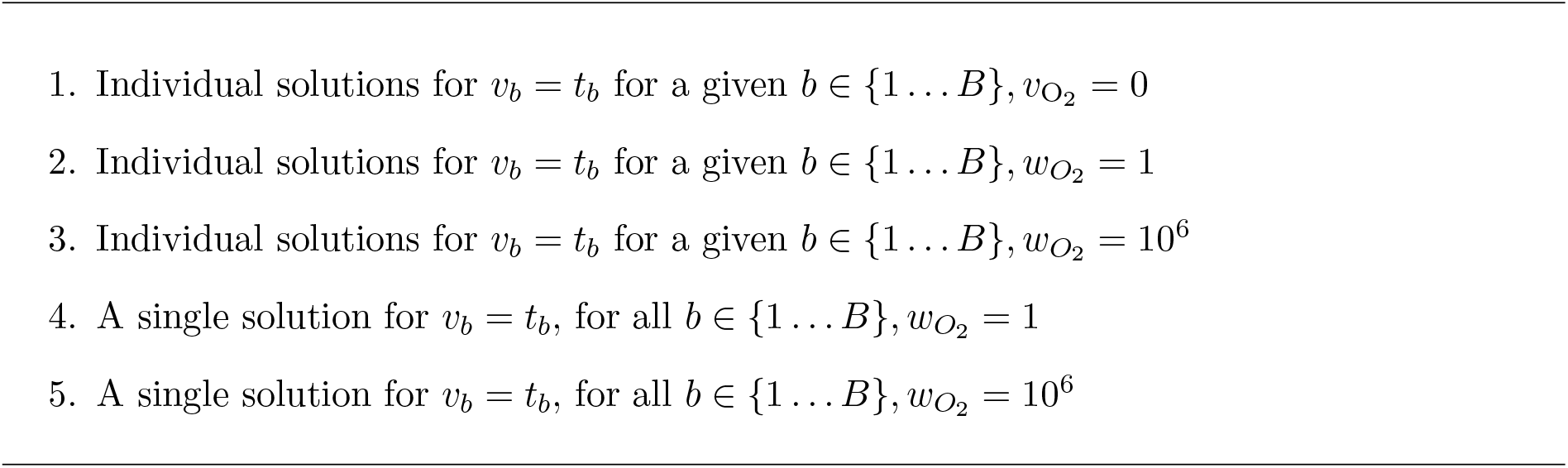
The five solution sets used to analyse the oxygen requirement of *C. jejuni*

In order to identify a solution accounting for all biomass components while minimising 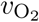, 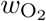, was set to an arbitrarily high value of 10^6^ and Eq. (1) re-solved. Thus five sets of flux distributions were obtained, as shown in Table 1.

**Table 1:** Modes of PLP production with or without a background demand for other biomass components and with or without O_2_ uptake penalty. The solutions for PLP production are single EMs.

| Penalty | +Biomass | Reactions | Total flux | O<sub>2</sub> uptake | Transporters |
| --- | --- | --- | --- | --- | --- |
| No | No | 46 | $4.6 \times 10^{-4}$ | $2.9 \times 10^{-5}$ | 10 |
| Yes | No | 47 | $5.0 \times 10^{-4}$ | $3.0 \times 10^{-6}$ | 10 |
| No | Yes | 140 | $1.1 \times 10^{-3}$ | $3.8 \times 10^{-5}$ | 15 |
| Yes | Yes | 136 | $1.3 \times 10^{-3}$ | $3.0 \times 10^{-6}$ | 16 |

### 2.3 Identifying EMs in a Steady-State Flux Vector

In order to identify individual EMs in LP solutions when all biomass components are produced simultaneously we developed an algorithm combining LP with null-space as follows.

The algorithm proceeds by iteratively removing one or more reactions at a time from an initial solution, **v**, and identifying an associated subsystem, **v**′, which either contains a single EM, or can be decomposed into a set of EMs. Then, **v**′ is subtracted from **v** and the process continues until **v** = **0**. We thereby obtain a matrix, **E**, whose column vectors consist of EMs, such that:

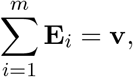

where *m* is the number of EMs. Note that, in this case, the EMs are not normalised in order for the magnitude of each **E**_*i*_ to reflect the contribution of that EM to **v**.

At each iteration, the reaction to be eliminated, **v**_targ_, is selected as the reaction with the smallest (absolute) flux in **v**. The elimination is achieved by obtaining a solution, **v**′, to the linear program:

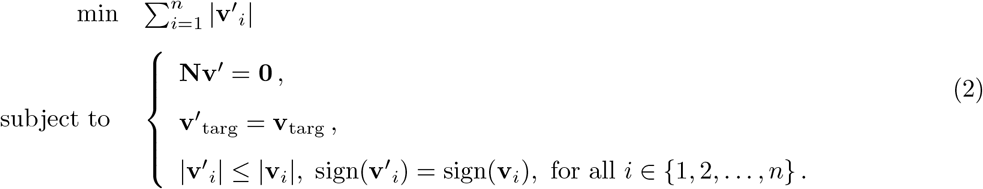

Given these definitions the algorithm can be described by the pseudo-code in presented in Algorithm 1.

#### Algorithm 1: LPEM - Using LP to decompose a steady-state flux vector, v, into a matrix of EMs, E

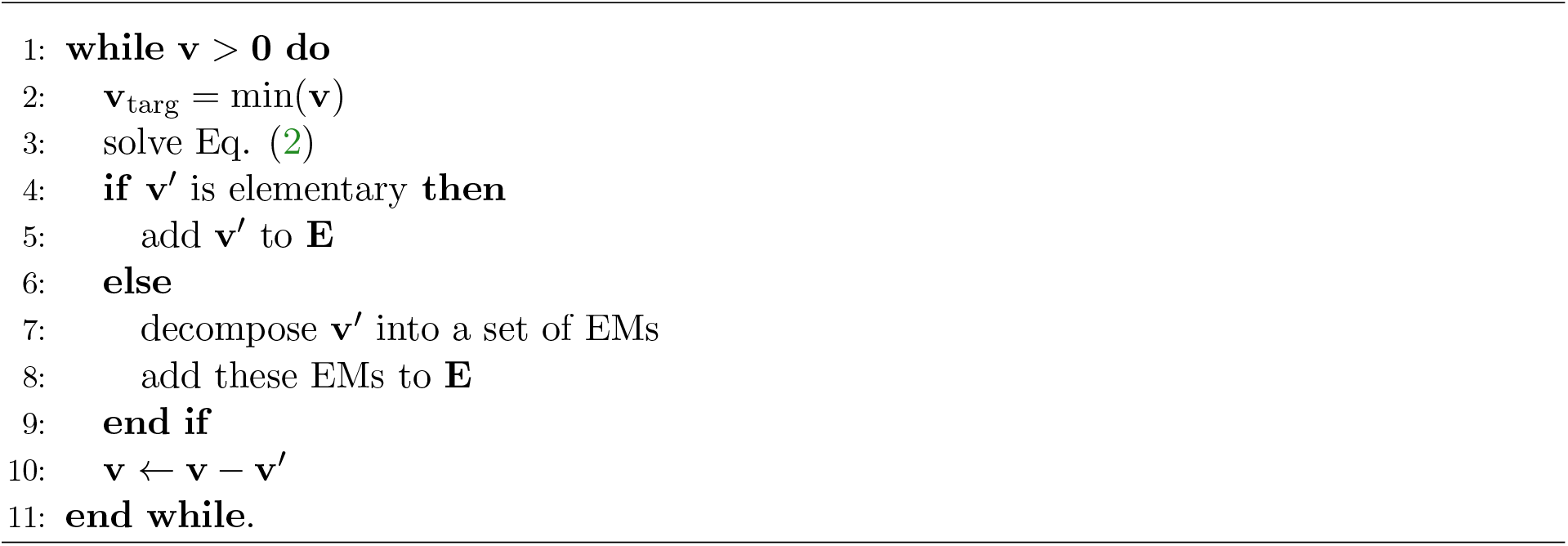

Verifying that **v**′ represents a single EM (step 4) is readily achieved by determining the dimension of null-space of the subsystem it represents: if it is 1, then **v**′ is an EM. If not, step 7 decomposes **v** into EMs using the algorithm by Schuster et al. [1999], and flux assigned to each EM as described in Poolman et al. [2004]. The constraints defined in Eq. (2) along with the subtraction in step 10 ensure that the number of non-zero elements in **v** decrease by at least one in each iteration of the algorithm, thereby guaranteeing completion.

## 3 Results

### 3.1 Model responses penalties and constraints described in Panel 1

No solution exists when attempting to solve Eq. (1) with the oxygen uptake constrained to zero, demonstrating an absolute requirement for oxygen to account for biomass production in this model. The results obtained under the different conditions described in Panel 1 were as follows:

1. **Individual biomass components, oxygen uptake set to zero** Solutions could be found for all individual biomass components, with the exception of pyridoxal 5′-phosphate (PLP), the active form of vitamin B6 and a co-substrate for many enzyme catalysed reactions (see discussion).
2. **Individual biomass components, with no constraint or penalty on oxygen uptake** With no constraints on oxygen uptake, the synthesis of 45 of the 51 biomass precursors utilised oxygen (including PLP synthesis), and all of these used proton translocating ATP synthase to satisfy some, or all, of their ATP demand; 43 (including PLP) utilised cytochrome oxidase and/or NADH oxidase to generate some or all of the necessary proton gradient to drive ATP synthesis.
3. **Individual biomass components, with imposed oxygen uptake penalty** With the oxygen uptake penalty (weighting factor 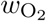 in Eq. (1)) set to 10^6^ none of the biomass precursors, with the exception of PLP, utilised oxygen. 43 of these (including PLP) utilised the electron transport chain for ATP generation with NO_3_ or NO_2_ acting as the terminal electron acceptor.
4. **Simultaneous production of all biomass components, with no oxygen uptake penalty** When Eq. (1) was solved to account for the simultaneous production of all biomass components, the resulting solution contained 324 reactions (excluding transport processes). The major carbon and nitrogen source was glutamine, accounting for 51% and 87% of total carbon and nitrogen uptake respectively. Most of the remaining carbon uptake was satisfied by the uptake of pyruvate (39%) with the rest of the carbon and nitrogen demand satisfied by serine, methionine and cysteine. The latter two amino acids satisfied the demand for sulphur. Excretion of carbon by-product was mainly in the form of carbon dioxide and carbonic acid (73%) with the remainder exported as acetate. Excess nitrogen was excreted in the form of NH_4_.
5. **Simultaneous production of all biomass components, with imposed oxygen uptake penalty** When the oxygen uptake penalty of 10^6^ was imposed on Eq. (1) with the demand for the synthesis of all biomass precursors, the resulting solution also contained 324 reactions of which 320 were common to the solution obtained with no imposed penalty. The fact that the two have the same number of reactions is assumed to be coincidence. All changes to the flux distribution (qualitative and quantitative) were associated with the redirection of the ETC from aerobic to anaerobic operation.

### 3.2 EMs of the production of biomass components

When solutions for individual biomass components were generated, no constraint was placed on the production of other biomass components, thus allowing the potential for the production of other components as by-products. Nonetheless, each solution resulted in the generation of the target product only, and represented a single EM (see discussion below). However, the total number of reactions utilised by all individual solutions was substantially greater than the number of reactions required to solve Eq. (1) for all components simultaneously (423 vs 397 and 434 vs 395 for penalised and unpenalised O_2_ uptake respectively). Therefore the solution obtained for the simultaneous production of all components is not simply the sum of the individual solutions for each component.

To further investigate this, the LPEM algorithm was used to identify EMs utilised for each biomass component in whole solutions (Panel 1.4 and 1.5). The sets of EMs thus calculated comprised a total of 62 EMs for both the penalised and unpenalised solutions. All of the unpenalised EMs utilised oxygen but only one (responsible for PLP synthesis) EM in the penalised set did so. Although the EM responsible for PLP synthesis with penalised O_2_ uptake was not the same as the equivalent LP solution, the rate of O_2_ uptake was the same for both.

The major difference between the EMs extracted from the whole solution and the individual LP solutions was that most EMs (all but one) generated more than one product and, conversely, most end products were generated by more than one EM.

### 3.3 EMs of PLP production

The solution for the synthesis of PLP in the absence of demand for other products is presented in Fig. 1 and summarised in Table 1. This is a superset of the the PLP synthesis pathway reported in metacyc (PYRIDOXSYN-PWY), which in turn is derived from the pathways proposed by Fitzpatrick et al. [2007] and described as the DXP-dependent and DXP-independent pathways. The pathway presented here is more complete as it balances all reactions starting from the external substrates glutamine and pyruvate, as well as generating necessary ATP and reductant. It is interesting to note that although both utilise the ETC for ATP generation, neither glycolysis (some reactions of glycolysis are present but run in the reverse, gluconeogenic, direction) nor the TCA cycle is used; sufficient reductant is generated by the oxidation of other substrates to satisfy the demand for reductant by the ETC. In fact, there is a slight excess of reductant generated which is then balanced by the reduction of external NO_3_ to NH_3_, which is in turn simply excreted (reactions R26 and R33 in Fig. 1).

**Figure 1:**
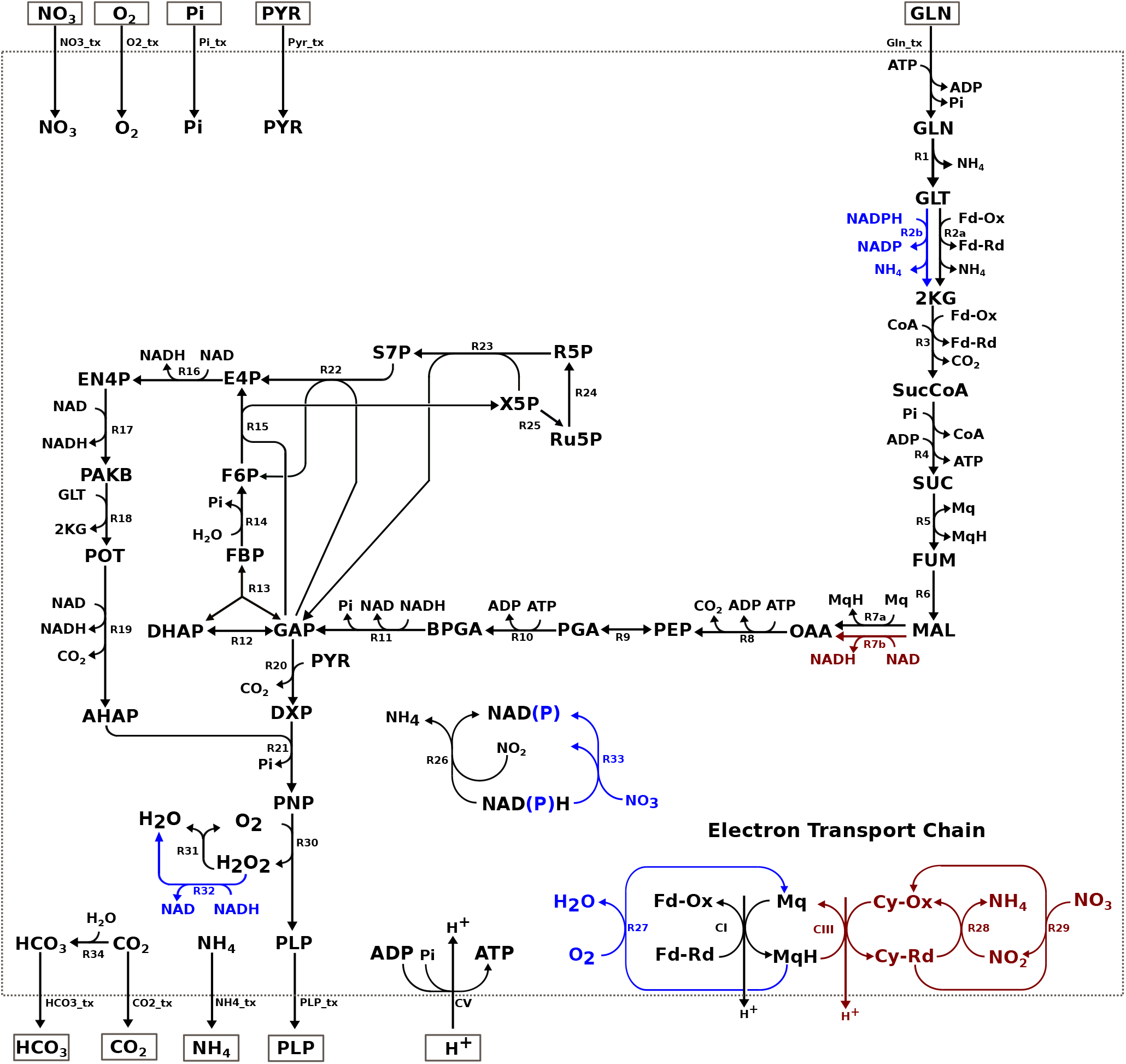
PLP synthesis in the *Campylobacter spp*. model with and without out a penalty on O_2_ uptake. Reactions in black are active under both the conditions. Reactions in blue are active when there is no penalty on O_2_ uptake and reactions in red are active only when the penalty is imposed. See key tables 1 and 2 for complete descriptions of reaction and metabolite abbreviations.

The EM obtained for the synthesis of PLP in the context of simultaneous synthesis of other biomass components, without imposing a penalty on O_2_ uptake is considerably more complex than the solution obtained for PLP as a single product (140 vs 46 reactions). However it does utilise the majority of the reactions depicted in Fig. 1, with the exception of those associated with oxidising excess reductant (R7a, R2a, R32, R33, R26 (NADP), R27). The other difference between this EM and the simple solution is that the EM also produces small amounts of other biomass components: DTTP, FAD, and valine.

A similar situation was found when comparing the EM producing PLP in the context of simultaneous synthesis of other biomass components, with the penalty on O_2_ imposed (136 vs 46 reactions), and contained 41 reactions in common with the simple solutions. Of the those that were absent, one was associated with the oxidation of excess reductant (nitrate reductase, R26 in Fig. 1), and two were associated with a move away from the use of membrane bound electron carriers to nicatinamides (malate oxidoreductase (R7a) and glutamate synthase (R2a) in Fig. 1). Somewhat unexpectedly, this EM did not utilise transaldolase (R22) or sedoheptulose 7-phosphate transketolase (R23) indicating that this EM has an alternative source of the pentose phosphate pathway intermediate, X5P. Again this EM also produced a number of other biomass precursors, in this case histidine, phosphatidyl-serine, FAD and valine.

### 3.4 Algorithm Performance

The LP solutions for production of all biomass components with free and penalised O_2_ uptake were comprised of 390 and 392 reactions respectively, the dimension of null-spaces of the subsystems defined by these solutions was 51 in each case. The LPEMs algorithm decomposed these two solutions into 52 and 55 EMs, remarkably close to the dimension of their respective null-spaces. Using a commodity desk-top PC with a 2.6 GHz AMD Ryzen processor the determination of the EMs for both solutions took about 70 seconds and required less than 1 GB of available memory.

## 4 Discussion and Conclusion

### 4.1 Micro-aerophilly in *C. jejuni*

The results presented above demonstrate that, in this model, O_2_ is essential for the synthesis of PLP, the biologically active form of vitamin B6. This is an enzyme bound co-factor for more than 140 reactions, mainly those involved in amino-acid metabolism and predominantly transferase and lyase reactions Eliot and Kirsch [2004], Percudani and Peracchi [2003, 2009]. Animals are unable to synthesise PLP, and it is therefore an essential vitamin. However, plants, fungi and bacteria are able to synthesize PLP via one of two reported routes: the DXP (deoxy-xylulose 5-phosphate) dependent and DXP independent pathway. In the DXP independent pathway, PLP is synthesised by single heterodimeric complex from glutamine, ribose 5-phosphate, and glyceraldehyde 3-phosphate. Organisms lacking this pathway, mainly the proteobacteria, utilise the DXP dependent pathway, commonly depicted as using erythrose 4-phosphate and glyceraldehyde 3-phosphate as the starting point. The pathway depicted in Fig. 1 is in fact a superset of the DXP dependent pathway.

Many other organisms which use the DXP dependent pathway have additional associated reactions and transporters, allowing the uptake of additional precursors as well as greater metabolic flexibility (Fig. 2), and in particular the potential to bypass the O_2_ dependent pyridoxine 5′-phosphate oxidase (R30 in Fig. 1 and 2) step Ito and Downs [2020], Sugimoto et al. [2017]. However, these have not been reported *C. jejuni* M1cam and therefore O_2_ is an absolute requirement for PLP synthesis.

**Figure 2:**
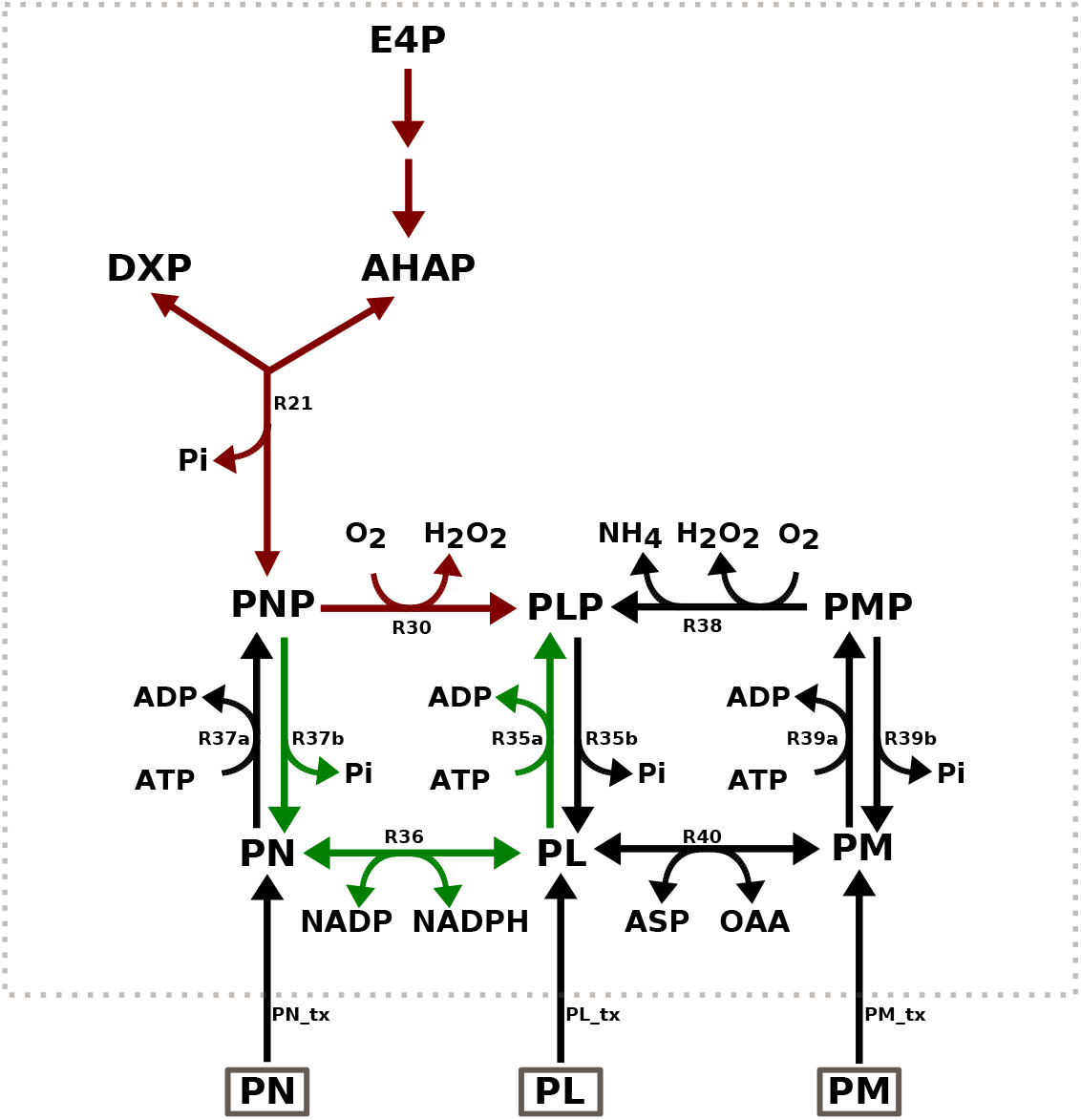
Reactions involved in PLP metabolism in *Escherichia coli*. Reactions in red are common to *C. jejuni*, those in green allow for the bypass of of the O_2_ dependent pyridoxine 5′-phosphate oxidase step (R30). *** Reactions labelled R35a/R37a/R39a are all catalysed by the same enzyme (similarly for reactions labelled R35b/R37b/R39b and R30/R38). See key tables 3 for a complete key.

### 4.2 Oxygen dependence of PLP synthesis

That the LP solutions of all biomass precursors individually, with free and restricted O_2_, were single EMs is unsurprising as the solutions of linear programs that contain only one non-zero flux constraint have been shown to always consist of a single EMs Maarleveld [2015]. What is more relevant is that this allows the unambiguous identification of PLP production as the reason for the absolute dependence on O_2_ for this model to account for growth, although this does not unambiguously identify which O_2_ consuming reactions are responsible for the dependence. The model contains a total of 27 reactions utilising O_2_ as a substrate, making it impractical to identify the essential reactions by a combinatorial search strategy. However the problem may be readily solved by using the technique of Enzyme Subsets analysis Pfeiffer et al. [1999] which identifies sets of reactions in a network which must carry flux in a fixed ratio in *any* steady-state. A corollary of this is that if any one reaction in a subset carries zero flux at a given steady state, then every other reaction must also carry zero flux. Determining the enzyme subsets of this model reveals that pyridoxine (pyridoxamine) 5′-phosphate oxidase (R30 in Fig. 1) is in the same subset as the PLP transporter and therefore it is the reaction responsible for the absolute dependency on O_2_ for the synthesis of PLP. It is interesting to note that, although the model contains the catalase reaction (R31 in Fig. 1), this cannot be used to generate internal O_2_ as a substrate for these reactions, as the generation of hydrogen peroxide itself must ultimately depend on an exogenous O_2_ source.

### 4.3 Sum of individual solutions compared to the decomposition of the whole solution

By using LP to identify pathways for precursor synthesis in isolation and applying the LPEM algorithm to an LP solution accounting for the simultaneous production of all precursors, two sets of results were obtained. Although it might be intuitively expected that the solution that accounts for all biomass precursors would be equivalent to the sum of the 51 pathways that produce each precursor individually, this was not the case: the summation of individual solutions required more reactions and greater total flux. This suggests that the optimal solution for the synthesis of a single product in isolation is not necessarily optimal in the presence of demand for additional products, and that therefore individual solutions must be interpreted with care in the context of a growing organism. The explanation for this appears to be that optimal individual solutions also generate by-products, that must then be further metabolised before they can be exported. However, when multiple products must be synthesised such by-products may be utilised for the synthesis of other products. For example, the solution for PLP synthesis in isolation (Fig. 1) generates excess reductant, which is then oxidised by the reduction of NO_3_ to NH_3_ which is subsequently exported. However, when there is a requirement to synthesise other products, these by-products become useful intermediates. A similar observation was originally made by Fell and Small [1986] in one of earliest papers describing the application of LP to metabolic networks.

### 4.4 Conclusion

The LPEM algorithm provides a relatively simple and computationally efficeint way to leverage the advantages of FBA and Elementary Modes Analysis. This has shown that the actual modes utilised by an organism *in vivo* may be rather more complicated than consideration of individual FBA solutions would suggest, but that these may nonetheless be more efficeint both in terms of the total number of reactions required and of the overall flux they carry. Applying the approaches described here suggests that the reason for micro-aerophilly in the pathogen *C. jejuni* is the dependence on oxygen for the production of PLP, although this may not be exclusive.

## Keys to figures

**Key 1.** Fig. 1 (Reactions)

| Label | Enzyme | Metacyc ID | EC number |
| --- | --- | --- | --- |
| R1 | glutaminase | GLUTAMIN-RXN | 3.5.1.38 |
| R2a | glutamate synthase | RXN-12878 | 1.4.7.1 |
| R2b | glutamate dehydrogenase | GLUTDEHYD-RXN | 1.4.1.4 |
| R3 | 2-oxoglutarate synthase | 2-OXOGLUTARATE-SYNTHASE-RXN | 1.2.7.3 |
| R4 | succinyl-CoA synthase | SUCCCOASYN-RXN | 6.2.1.5 |
| R5 | succinate dehydrogenase | SUCCINATE-DEHYDROGENASE-MENAQUINONE-RXN | 1.3.5.1 |
| R6 | fumerase | FUMHYDR-RXN | 4.2.1.2 |
| R7a | malate oxidoreductase (quinone) | RXNI-3 | 1.1.5.4 |
| R7b | malate dehydrogenase | MALATE-DEH-RXN | 1.1.1.37 |
| R8 | phosphoenolpyruvate carboxykinase | PEPCARBOXYKIN-RXN | 4.1.1.49 |
| R9 | enolase | 2PGADEHYDRAT-RXN | 4.2.1.11 |
| R10 | phosphoglycerate kinase | PHOSGLYPHOS-RXN | 2.7.2.3 |
| R11 | glyceraldehyde 3-phosphate dehydrogenase | GAPOXNPHOSPHN-RXN | 1.2.1.12 |
| R12 | triose-phosphate isomerase | TRIOSEPISOMERIZATION-RXN | 5.3.1.1 |
| R13 | fructose-bisphosphate aldolase | F16ALDOLASE-RXN | 4.1.2.13 |
| R14 | fructose 1,6-bisphosphatase | F16BDEPHOS-RXN | 3.1.3.11 |
| R15 | fructofuranose 6-phosphate transketolase | 2TRANSKETO-RXN | 2.2.1.1 |
| R16 | erythrose 4-phosphate dehydrogenase | ERYTH4PDEHYDROG-RXN | 1.2.1.72 |
| R17 | erythronate 4-phosphate dehydrogenase | ERYTHRON4PDEHYDROG-RXN | 1.1.1.290 |
| R18 | phosphohydroxythreonine aminotransferase | PSERTRANSAMPYR-RXN | 2.6.1.52 |
| R19 | hydroxythreonine 4-phosphate dehydrogenase | RXN-13179 | 1.1.1.262 |
| R20 | deoxy-xylulose 5-phosphate synthase | DXS-RXN | 2.2.1.7 |
| R21 | pyridoxine 5'-phosphate synthase | PDXJ-RXN | 2.6.99.2 |
| R22 | transaldolase | TRANSALDOL-RXN | 2.2.1.2 |
| R23 | seduloheptulose 7-phosphate transketolase | 1TRANSKETO-RXN | 2.2.1.1 |
| R24 | ribose 5-phosphate isomerase | RIB5PISOM-RXN | 5.3.1.6 |
| R25 | ribulose phosphate 3-epimerase | RIBULP3EPIM-RXN | 5.1.3.1 |
| R26 | nitrite reductase (NAD) | RXN0-6377 | 1.7.1.4 |
| R27 | oxygen reductase (cytochrome) | RXN0-5266 | 7.1.1.7 |
| R28 | nitrite reductase (cytochrome) | 1.7.2.2-RXN | 1.7.2.2 |
| R29 | nitrate reductase (cytochrome) | NITRATE-REDUCTASE-CYTOCHROME-RXN | 1.9.6.1 |
| R30 | pyridoxine 5'-phosphate oxidase | PNPOXI-RXN | 1.4.3.5 |
| R31 | catalase | CATAL-RXN | 1.11.1.6 |
| R32 | NADH peroxidase | NADH-PEROXIDASE-RXN | 1.11.1.1 |
| R33 | nitrate reductase (NAD) | NITRATE-REDUCTASE-NADPORNOPH-RXN | 1.7.99.4 |
| R34 | carbonic anhydrase | RXN0-5224 | 4.2.1.1 |
| CV | proton translocating ATP synthase | ATPSYN-RXN | 7.1.2.2 |

**Key 2.**
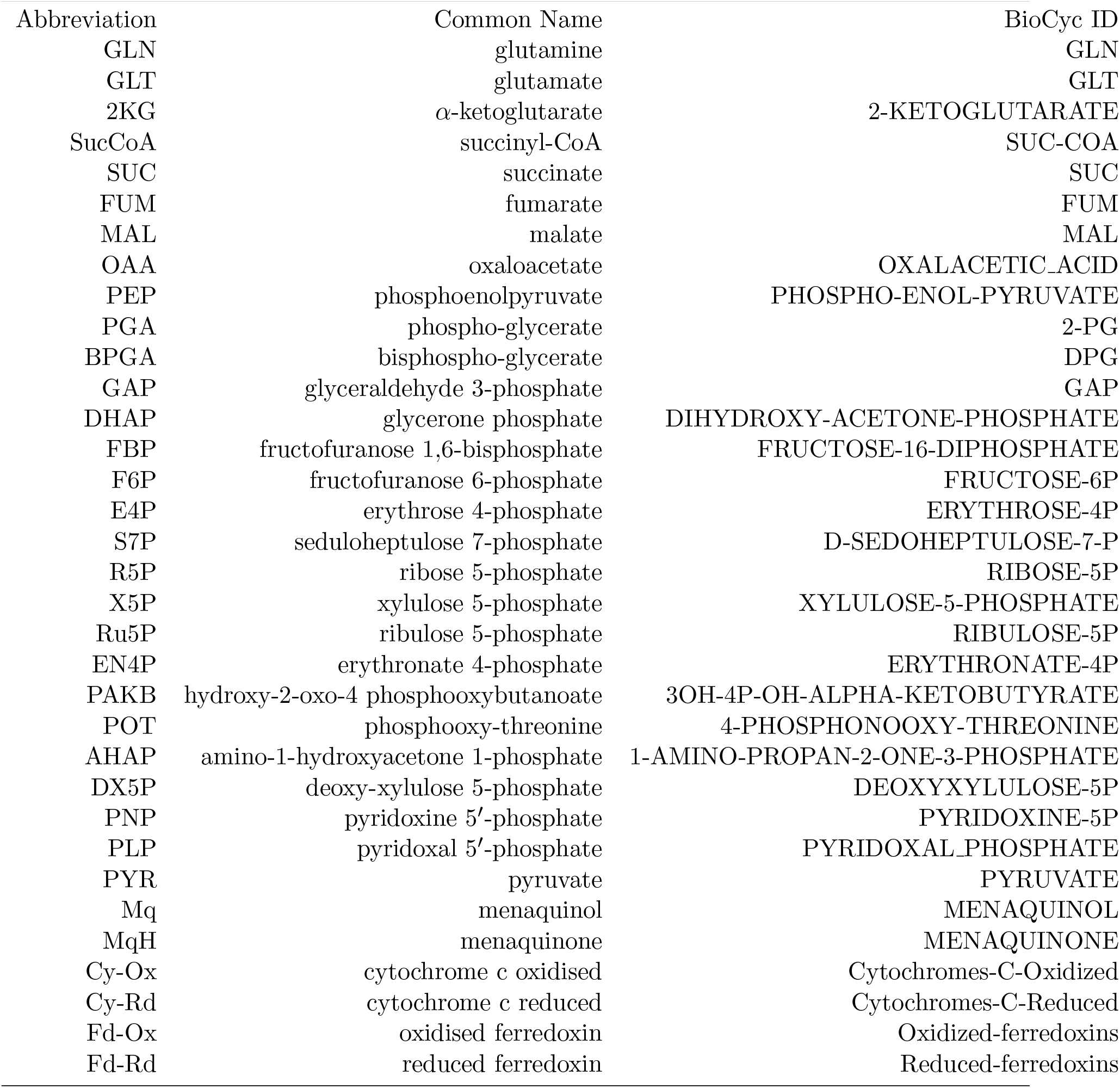
Fig. 1 (Metabolites)

**Key 2** Fig. 1 (Metabolites)
| Abbreviation | Common Name | BioCyc ID |
| --- | --- | --- |
| GLN | glutamine | GLN |
| GLT | glutamate | GLT |
| 2KG | α-ketoglutarate | 2-KETOGLUTARATE |
| SucCoA | succinyl-CoA | SUC-COA |
| SUC | succinate | SUC |
| FUM | fumarate | FUM |
| MAL | malate | MAL |
| OAA | oxaloacetate | OXALACETIC_ACID |
| PEP | phosphoenolpyruvate | PHOSPHO-ENOL-PYRUVATE |
| PGA | phospho-glycerate | 2-PG |
| BPGA | bisphospho-glycerate | DPG |
| GAP | glyceraldehyde 3-phosphate | GAP |
| DHAP | glycerone phosphate | DIHYDROXY-ACETONE-PHOSPHATE |
| FBP | fructofuranose 1,6-bisphosphate | FRUCTOSE-16-DIPHOSPHATE |
| F6P | fructofuranose 6-phosphate | FRUCTOSE-6P |
| E4P | erythrose 4-phosphate | ERYTHROSE-4P |
| S7P | seduloheptulose 7-phosphate | D-SEDOHEPTULOSE-7-P |
| R5P | ribose 5-phosphate | RIBOSE-5P |
| X5P | xylulose 5-phosphate | XYLULOSE-5-PHOSPHATE |
| Ru5P | ribulose 5-phosphate | RIBULOSE-5P |
| EN4P | erythronate 4-phosphate | ERYTHRONATE-4P |
| PAKB | hydroxy-2-oxo-4 phosphooxybutanoate | 3OH-4P-OH-ALPHA-KETOBUTYRATE |
| POT | phosphooxy-threonine | 4-PHOSPHONOOXY-THREONINE |
| AHAP | amino-1-hydroxyacetone 1-phosphate | 1-AMINO-PROPAN-2-ONE-3-PHOSPHATE |
| DX5P | deoxy-xylulose 5-phosphate | DEOXYXYLULOSE-5P |
| PNP | pyridoxine 5'-phosphate | PYRIDOXINE-5P |
| PLP | pyridoxal 5'-phosphate | PYRIDOXAL_PHOSPHATE |
| PYR | pyruvate | PYRUVATE |
| Mq | menaquinol | MENAQUINOL |
| MqH | menaquinone | MENAQUINONE |
| Cy-Ox | cytochrome c oxidised | Cytochromes-C-Oxidized |
| Cy-Rd | cytochrome c reduced | Cytochromes-C-Reduced |
| Fd-Ox | oxidised ferredoxin | Oxidized-ferredoxins |
| Fd-Rd | reduced ferredoxin | Reduced-ferredoxins |

**Key 3.**
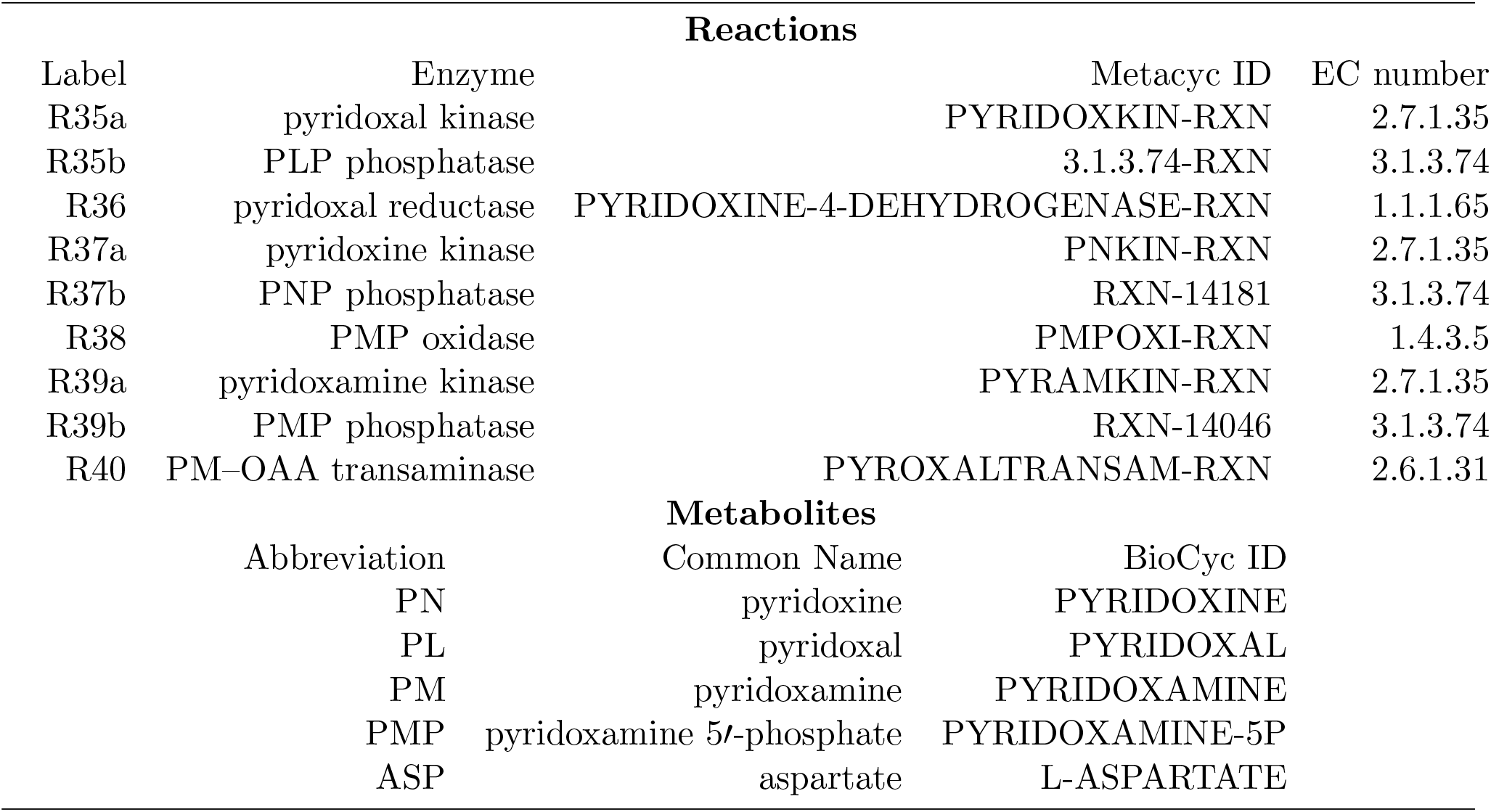
Fig. 2

## Acknowledgements

**YS** The research work disclosed in this publication is partially funded by the *Tertiary Education Scholarships Scheme* (Malta) and Oxford Brookes University (*Nigel Groome Studentship*)

**DS** Is funded by the UK Biotechnology and Biological Sciences Research Council (BBSRC), through the Institute Strategic Research Programme (ISP) BB/R012504/1 Microbes in the Food Chain and its constituent project Microbial Survival in the Food Chain BB-S/E/F/000PR10349.

**MP** Is funded by Oxford Brookes University and the European Union’s Horizon 2020 research and innovation programme under the Marie Sklodowska-Curie grant agreement number 956154 (INNOTARGETS).

## Footnotes

1 https://www.gov.uk/government/publications/campylobacter-infection-annual-data, accessed December 2022.

## References

Vicente Acuña, Flavio Chierichetti, Vincent Lacroix, Alberto Marchetti-Spaccamela, Marie France Sagot, and Leen Stougie. Modes and cuts in metabolic networks: Complexity and algorithms. BioSystems, 95(1):51–60, 2009. ISSN 03032647. doi: 10.1016/j.biosystems.2008.06.015.

Lindsey Albenberg, Tatiana V Esipova, Colleen P Judge, Kyle Bittinger, Jun Chen, Alice Laughlin, Stephanie Grunberg, Robert N Baldassano, James D Lewis, Hongzhe Li, Stephen R Thom, Frederic D Bushman, Sergei A Vinogradov, and Gary D Wu. Correlation between intraluminal oxygen gradient and radial partitioning of intestinal microbiota. Gastroenterology, 147(5): 1055–63.e8, November 2014. doi: 10.1053/j.gastro.2014.07.020.

A. Bernalier-Donadille. Fermentative metabolism by the human gut microbiota. Gastroentérologie Clinique et Biologique, 34:S16–S22, 2010. ISSN 0399-8320. doi: 10.1016/S0399-8320(10)70016-6.

Nicolas Daniel, Nuria Casadevall, Pei Sun, Daniel Sugden, and Vanna Aldin. The burden of foodborne disease in the UK 2018. https://www.food.gov.uk/sites/default/files/media/document/the-burden-of-foodborne-disease-in-the-uk_0.pdf, 2020.

Stefan P. W. de Vries, Srishti Gupta, Abiyad Baig, Joanna Lapos Heureux, Elsa Pont, Dominika P. Wolanska, Duncan J. Maskell, and Andrew J. Grant. Motility defects in Campylobacter jejuni defined gene deletion mutants caused by second-site mutations. Microbiology, 161(12):2316–2327, 2015. ISSN 1465-2080. doi: 10.1099/mic.0.000184.

Andrew C. Eliot and Jack F. Kirsch. Pyridoxal phosphate enzymes: Mechanistic, structural, and evolutionary considerations. Annual Review of Biochemistry, 73(1):383–415, 2004. doi: 10.1146/annurev.biochem.73.011303.074021.

David a Fell and J R Small. Fat synthesis in adipose tissue. The Biochemical journal, 238(3): 781–786, 1986. ISSN 02646021. URL http://www.pubmedcentral.nih.gov/articlerender.fcgi?artid=1147204&tool=pmcentrez&rendertype=abstract.

Teresa B. Fitzpatrick, Nikolaus Amrhein, Barbara Kappes, Peter Macheroux, Ivo Tews, and Thomas Raschle. Two independent routes of de novo vitamin b6 biosynthesis: Not that different after all. Biochemical Journal, 407(1):1–13, 2007. doi: 10.1042/bj20070765.

RDM Hadden, NA Gregson, R Gold, KJ Smith, and RAC Hughes. Accumulation of immunoglobulin across the ‘blood-nerve barrier’ in spinal roots in adoptive transfer experimental autoimmune neuritis. Neuropathology and Applied Neurobiology, 28(6):489–497, 2002. ISSN 0305-1846. doi: 10.1046/j.1365-2990.2002.00421.x.

Hassan B. Hartman, David A. Fell, Sergio Rossell, Peter Ruhdal Jensen, Martin J. Woodward, Lotte Thorndahl, Lotte Jelsbak, John Elmerdahl Olsen, Anu Raghunathan, Simon Daefler, and Mark G. Poolman. Identification of potential drug targets in salmonella enterica sv. typhimurium using metabolic modelling and experimental validation. Microbiology, 160(6): 1252–1266, 2014. doi: 10.1099/mic.0.076091-0.

Dirk Hofreuter. Defining the metabolic requirements for the growth and colonization capacity of Campylobacter jejuni. Frontiers in Cellular and Infection Microbiology, 4, 2014. ISSN 2235-2988. doi: 10.3389/fcimb.2014.00137.

Siu Hung, Joshua Chan, and Ping Ji. Decomposing flux distributions into elementary flux modes in genome-scale metabolic networks. Bioinformatics, 27(16):2256–2262, 2011. ISSN 14602059. doi: 10.1093/bioinformatics/btr367.

Tomokazu Ito and Diana M. Downs. Pyridoxal reductase, pdxi, is critical for salvage of pyridoxal in Escherichia coli. Journal of Bacteriology, 202(12):e00056–20, 2020. doi: 10.1128/JB.00056-20. URL https://journals.asm.org/doi/abs/10.1128/JB.00056-20.

Raphael M. Jungers, Francisca Zamorano, Vincent D. Blondel, Alain Vande Wouwer, and Georges Bastin. Fast computation of minimal elementary decompositions of metabolic flux vectors. Automatica, 47(6):1255–1259, 2011. ISSN 00051098. doi: 10.1016/j.automatica.2011.01.011.

Nadeem O. Kaakoush, William G. Miller, Hilde De Reuse, and George L. Mendz. Oxygen requirement and tolerance of Campylobacter jejuni. Research in Microbiology, 158(8-9):644–650, OCT-NOV 2007. ISSN 0923-2508. doi: 10.1016/j.resmic.2007.07.009.

David J. Kelly. Complexity and versatility in the physiology and metabolism of Campylobacter jejuni. In Campylobacter, Third Edition, pages 41–61. American Society of Microbiology, 2008. doi: 10.1128/9781555815554.ch3.

S. Klamt and E.D. Gilles. Minimal cut sets in biochemical reaction networks. Bioinformatics, 20(2):226–234, 2004. doi: 10.1093/bioinformatics/btg395.

Timo Maarleveld. Fluxes and Fluctuations in Biochemical Models. PhD thesis, Vrije Universiteit Amsterdam, 2015.

Noah Adam Mesfin. Structural metabolic modelling of the acetogen Acetobacterium woodii. PhD thesis, Oxford Brookes University, 2020.

Hildur Æsa Oddsdóttir, Erika Hagrot, Véronique Chotteau, and Anders Forsgren. On dynamically generating relevant elementary flux modes in a metabolic network using optimization. Journal of Mathematical Biology, 71(4):903–920, 2015. ISSN 14321416. doi: 10.1007/s00285-014-0844-1.

Jeffrey D. Orth, Ines Thiele, and Bernhard O. Palsson. What is flux balance analysis? Nature Biotechnology, 28(3):245–248, 2010. ISSN 10870156. doi: 10.1038/nbt.1614.

Julianne Parkhill, B Wren, K Mungall, J Ketley, Carol Churcher, Daryl Basham, T Chillingworth, R Davies, T Feltwell, S Holroyd, K Jagels, Andrey Karlyshev, S Moule, Mark Pallen, C Penn, MA Quail, MA Rajandream, K Rutherford, Arnoud van Vliet, and B Barrell. The genome sequence of the food-borne pathogen Campylobacter jejuni reveals hypervariable sequences. Nature, 403:665–8, 03 2000. doi: 10.1038/35001088.

Riccardo Percudani and Alessio Peracchi. A genomic overview of pyridoxal-phosphate-dependent enzymes. EMBO reports, 4(9):850–854, 2003. doi: 10.1038/sj.embor.embor914.

Riccardo Percudani and Alessio Peracchi. The b6 database: A tool for the description and classification of vitamin b6-dependent enzymatic activities and of the corresponding protein families. BMC bioinformatics, 10:273, 10 2009. doi: 10.1186/1471-2105-10-273.

T. Pfeiffer, I. Sanchez-Valdenebro, J.C. Nuno, F. Montero, and S. Schuster. Metatool: for studying metabolic networks. Bioinformatics, 15(3):251–257, 1999. doi: 10.1093/bioinformatics/15.3.251.

M. G. Poolman. Scrumpy: metabolic modelling with python. Systems biology, 153(5):375–378, 2006. doi: 10.1049/ip-syb:20060010.

M. G. Poolman, K. V. Venkatesh, M. K. Pidcock, and D. A. Fell. A method for the determination of flux in elementary modes, and its application to Lactobacillus rhamnosus. Biotechnol. Bioeng., 88(5):601–612, 2004. doi: 10.1002/bit.20273.

Mark G. Poolman. Scrumpy. gitlab.com/MarkPoolman/scrumpy, 2020.

Kate O. Poropatich, Christa L. Fischer Walker, and Robert E. Black. Quantifying the association between Campylobacter infection and Guillain-Barre syndrome: A systematic review. Journal of Health Population and Nutrition, 28(6):545–552, DEC 2010. ISSN 1606-0997. doi: 10.3329/jhpn.v28i6.6602.

Christophe H. Schilling, Jeremy S. Edwards, David Letscher, and Bernhard Ø. Palsson. Combining pathway analysis with flux balance analysis for the comprehensive study of metabolic systems. Biotechnol Bioeng., 71(4):286–306, 2000.

S. Schuster and C. Hilgetag. On elementary flux modes in biochemical systems at steady state. J.Biol.Syst., 2:165–182, 1994. doi: 10.1142/S0218339094000131.

S. Schuster, T. Dandekar, and D.A. Fell. Detection of elementary flux modes in biochemical networks: a promising tool for pathway analysis and metabolic engineering. Trends. Biotech., 17(2):53–60, 1999. doi: 10.1016/s0167-7799(98)01290-6.

S. Schuster, D.A. Fell, and T. Dandekar. A general definition of metabolic pathways useful for systematic organization and analysis of complex metabolic networks. Nature Biotech., 18: 326–332, 2000. doi: 10.1038/73786.

J.-M. Schwartz and M. Kanehisa. A quadratic programming approach for decomposing steadystate metabolic flux distributions onto elementary modes. Bioinformatics, 21(Suppl 2):ii204–ii205, 2005. ISSN 1460-2059. doi: 10.1093/bioinformatics/bti1132. URL http://dx.doi.org/10.1093/bioinformatics/bti1132.

Michael J. Sellars, Stephen J. Hall, and David J. Kelly. Growth of Campylobacter jejuni supported by respiration of fumarate, nitrate, nitrite, trimethylamine-n-oxide, or dimethyl sulfoxide requires oxygen. Journal of Bacteriology, 184(15):4187–4196, 2002. ISSN 0021-9193. doi: 10.1128/JB.184.15.4187-4196.2002.

Martin Stahl, James Butcher, and Alain Stintzi. Nutrient acquisition and metabolism by Campylobacter jejuni. Frontiers in Cellular and Infection Microbiology, 2, 2012. ISSN 2235-2988. doi: 10.3389/fcimb.2012.00005.

Emily Stoakes, George M. Savva, Ruby Coates, Noemi Tejera, Mark G. Poolman, Andrew J. Grant, John Wain, and Dipali Singh. Substrate utilisation and energy metabolism in non-growing Campylobacter jejuni M1cam. Microorganisms, 10(7), 2022. ISSN 2076-2607. doi: 10.3390/microorganisms10071355.

Ryota Sugimoto, Natsumi Saito, Tomohiro Shimada, and Kan Tanaka. Identification of ybha as the pyridoxal 5’-phosphate (plp) phosphatase in Escherichia coli: Importance of plp homeostasis on the bacterial growth. The Journal of General and Applied Microbiology, 63(6): 362–368, 2017. doi: 10.2323/jgam.2017.02.008.

Noemi Tejera, Lisa Crossman, Bruce Pearson, Emily Stoakes, Fauzy Nasher, Bilal Djeghout, Mark Poolman, John Wain, and Dipali Singh. Genome-scale metabolic model driven design of a defined medium for Campylobacter jejuni M1cam. Frontiers in Microbiology, 11, JUN 19 2020. ISSN 1664-302X. doi: 10.3389/fmicb.2020.01072.

Anne-Xander van der Stel and Marc M. S. M. Wösten. Regulation of respiratory pathways in campylobacterota: A review. Frontiers in Microbiology, 10, 2019. ISSN 1664-302X. doi: 10.3389/fmicb.2019.01719.

Anne-Xander van der Stel, Fred C. Boogerd, Steven Huynh, Craig T. Parker, Linda van Dijk, Jos P. M. van Putten, and Marc M. S. M. Wösten. Generation of the membrane potential and its impact on the motility, atp production and growth in Campylobacter jejuni. Molecular Microbiology, 105(4):637–651, 2017. doi: 10.1111/mmi.13723.

Sariqa Wagley, Jane Newcombe, Emma Laing, Emmanuel Yusuf, Christine M. Sambles, David J. Studholme, Roberto M. La Ragione, Richard W. Titball, and Olivia L. Champion. Differences in carbon source utilisation distinguish Campylobacter jejuni from Campylobacter coli. BMC Microbiology, 14, 2014. ISSN 1471-2180. doi: 10.1186/s12866-014-0262-y.

Dilan R. Weerakoon, Nathan J. Borden, Carrie M. Goodson, Jesse Grimes, and Jonathan W. Olson. The role of respiratory donor enzymes in Campylobacter jejuni host colonization and physiology. Microbial Pathogenesis, 47(1):8–15, 2009. ISSN 0882-4010. doi: 10.1016/j.micpath.2009.04.009.

Leon Zheng, Caleb J Kelly, and Sean P Colgan. Physiologic hypoxia and oxygen homeostasis in the healthy intestine. A Review in the Theme: Cellular Responses to Hypoxia. Am J Physiol Cell Physiol, 309(6):C350–60, 2015. doi: 10.1152/ajpcell.00191.2015.

